# Threatened terrestrial vertebrates are exposed to human pressures across 94% of Europe’s protected land

**DOI:** 10.1101/2020.08.25.266379

**Authors:** Dobrochna M. Delsen, W. Daniel Kissling, James R. Allan

## Abstract

Protected areas (PAs) are the last refuges for wild biodiversity, yet human pressures (or threats) are increasingly prevalent within their boundaries. Human pressures have the potential to negatively impact species and undermine their conservation, but their overlap with sensitive threatened species in PAs remains rarely quantified. Here, we analyse the co-occurrence of nineteen threatening human activities within the distributions of 146 threatened terrestrial vertebrates in the European Union (EU), accounting for species-specific sensitivities to each pressure and thereby mapping potential human impacts on species within EU PAs. We find that human pressures extend across > 1.022 million km^2^ (94.5%) of EU protected land, with potential negative impacts on threatened species across 1.015 million km^2^ (93.8%). A total of 122 out of 146 species (84%) have > 50% of their EU protected ranges potentially impacted, and 83 out of 146 species (57%) more than 90% of their protected range. More species have a smaller proportion of their protected range potentially impacted in Natura2000 sites than in non-Natura2000 sites, and the same is true for species in nature reserves and wilderness areas compared to less strictly managed PAs. Our results show that threatened species in Europe’s PAs are exposed to immense human pressures, and suggest that areas designated for species conservation are ineffective for halting biodiversity decline. We recommend that the EU Biodiversity Strategy develops and enforces a comprehensive PA threat management program to reduce the negative impacts of human activities on wildlife in European protected lands.

## 1. Introduction

In May 2020 the European Commission adopted a new biodiversity strategy with an ambitious plan to improve the state of Europe’s wildlife and habitats by 2030^1^. The main elements of this strategy include the expansion of protected areas (PAs) and restoration of degraded ecosystems, which align with targets in the Convention on Biological Diversity’s (CBD’s) post-2020 global biodiversity framework^1,2^. PAs are the main tool for biodiversity conservation worldwide and are effective when managed well^3^, but many are failing to meet their conservation potential^4^. One reason for this is that threats to biodiversity are prevalent in PAs^5–7^. For example, one-third of global protected land is exposed to intense human pressures^8^. The challenge of threats in protected areas is especially true in Europe, a continent with a long history of humans modifying the environment^9^.

The European Union (EU) has made efforts to conserve its most iconic and endangered species and habitats under the legal frameworks of the 1979 Birds Directive^10^ and the 1992 Habitats Directive^11^. These directives led to the creation of the Natura2000 coordinated PA network encompassing over 27,300 sites^12^, which is one of the largest PA networks in the world and the cornerstone of the EU biodiversity conservation strategy. PAs can be classified under management categories with varying levels of human activities, which can range from strict to more flexible protection from human use^13^. Natura2000 sites do not necessarily restrict human activity, and instead encourage humans and nature to co-exist sustainably. It is therefore not surprising that the majority of PAs in Europe contain human activities^8,14,15^, which could potentially compromise their goal of conserving Europe’s threatened species and habitats if human activities constitute a threat for protected species^16^. Yet, comprehensive assessments of human pressures within European protected lands and their connection to threatened species conservation are largely missing.

The presence of human pressures (often called threats) — defined as human activities or land uses with the potential to damage nature^17^ — does not always translate into a negative impact on biodiversity. This is because a pressure must co-occur with a species that is sensitive to that pressure for a negative impact to be realised^18^. An example of an activity that demonstrates this is human recreation such as hiking. Recreational hiking can have a negative impact on sensitive species^19,20^ but may be of little concern for other species. For instance, large predators can change their behaviour and habitat use as a response to recreational hiking, moving away from hiking trails due to anthropogenic fear^21^. Conversely, some herbivore and small mammal species are less sensitive to human intrusion and are not harmed by recreational hiking, or may even benefit from using areas of higher human activity to benefit from reduced predation risk^21,22^. Similarly to recreational activities, residential development and urbanization can have a substantial negative impact on many species due to habitat loss and ecosystem degradation^23^, but this may also benefit some species, e.g. bats that roost in attics or cellars of urban environments^24^. Accounting for species-specific sensitivity to threats is therefore crucial when studying the impacts of human pressures on species in PAs.

Previous efforts to analyse threats in PAs have mapped the distribution of human pressures within PA boundaries^5,7,8,25^, but did not account for the spatial overlap between threats and species geographic distributions^26^, or whether the species are sensitive to those threats^18^. Therefore, the causal link these studies make between human pressures and potential negative impacts on species is somewhat ambiguous, and means that impacts on biodiversity could be overestimated because pressures are assumed to impact all species. Indeed, numerous species can survive or even thrive in areas under some form of human activity^27,28^. Therefore, making the distinction between acceptable non-damaging human activities in PAs, and activities which negatively impact species, is crucial for PA managers to regulate human presence in a way that allows humans to still access resources and utilise the land while simultaneously guaranteeing that species are effectively conserved. Without this information there is a risk that either all human activities get unnecessarily prohibited from PAs, which could have social impacts on local communities and create conflicts^29^, or that harmful human activities and land uses are allowed within PAs to the detriment of the species they are meant to safeguard^30^.

Here, we address this gap by carrying out a comprehensive spatial analysis of human pressures within EU PAs and how they overlap with the distributions of threatened terrestrial vertebrate species while accounting for species-specific threat-sensitivities^18^. This method allows us to answer four timely questions for EU biodiversity conservation; 1) how are species-specific pressures spatially distributed across Europe, 2) to what extent do human pressures coincide with the distribution of sensitive threatened species in EU PAs, 3) are threatened species potentially less impacted in Natura2000 sites than in PAs outside Natura2000, or when species receive special policy attention under the EU Birds or Habitats Directive, and 4) how does the extent and type of human pressure on threatened species differ between PA management classes of varying management strictness. By answering those questions our results provide important information for supporting the new EU biodiversity strategy and ensuring effective conservation of European biodiversity in protected areas.

## 2. Methods

### 2.1 Protected areas in the European Union

PAs in the EU consist of nationally designated sites, and Special Areas of Conservation (SAC) and Special Protection Areas (SPA) established under Annex I of the 1979 Birds Directive^10^ and Annex I and II of the 1992 Habitats Directive^11^. SAC and SPA are selected on scientific grounds and consists of core areas meant to protect species and habitat types listed in the previously mentioned Annexes. SAC and SPA make up the Natura2000 network. The International Union for Conservation of Nature (IUCN) defines 6 management categories for PAs which are strict nature reserves (Ia), wilderness areas (Ib), national parks (II), natural monument/feature (III), habitat/species management area (IV), protected landscape/seascape (V) and PAs with sustainable use of natural resources (VI). Different human activities are allowed under different PA management categories; however, the primary aim of all PAs with an IUCN category is first and foremost biodiversity conservation^13^.

### 2.2 Collecting spatial data

#### 2.2.1 Protected areas and the Natura2000 network

We obtained spatial data on terrestrial nationally designated protected sites as well as information on their IUCN management class from the January 2020 World Database on PAs^31^. Following Jones *et al.* (2018)^8^ we only included PAs having a status of ‘designated’, ‘established’ or ‘inscribed’ and that were not designated as UNESCO Man and Biosphere Reserves. Additional spatial data on PAs from the terrestrial Natura2000 network (2018) were obtained from the European Environment Agency^32^. To reduce computational burden, we removed contiguous PAs < 1 km^2^ from the PA and Natura2000 data. We then merged the PAs from the World Database and the Natura2000 network into one layer and overlaid it with a 3×3 km grid. To avoid excessive grid cell fragments along edges, we removed any grid cells with an area < 0.09 km2 (this resulted in the exclusion of only 0.16% of total area).

Grid cells were assigned to be part of the Natura2000 network when a PA under the Natura2000 network overlapped > 50% with the area of the grid cell. The same threshold of > 50% area overlap was used for assigning a PA management class per grid cell. If multiple PA management classes had > 50% area overlap within one grid cell, we assigned the PA management class with highest naturalness (related to management strictness, ecosystem structure and human activity) to the grid cell (Ia > Ib > II > III > IV > VI > V > N)^33^.

#### 2.2.2 Threatened species ranges

We focussed on threatened terrestrial vertebrates (amphibians, reptiles, birds and mammals) with available distribution data (expert range maps) and IUCN threat assessments. Amphibian and mammal distribution data were obtained from the IUCN RedList^23^, distribution data on birds from BirdLife International^34^ and distribution data on reptiles from Roll *et al.* (2017)^35^. Threat assessment data were derived for all included vertebrates from the IUCN RedList^23^. We focussed only on species that were listed as near threatened, vulnerable, endangered or critically endangered on a global level since their species-specific threats have been comprehensively assessed. Following Allan *et al.* (2019)^18^, we used the extant geographic ranges of native or reintroduced species that (completely or partially) fall within the borders of protected land in the European Union. We excluded possibly extant ranges and parts of species distributions with uncertain or no current records of species presences, as well as introduced and vagrant species, and species with unknown origin. We did not filter species ranges based on seasonality (e.g. breeding ranges of birds) and instead included both winter and summer ranges. Based on all our criteria the distributions of 28 amphibian, 29 reptile, 59 bird and 30 mammal species qualified for the analyses. We considered a species to be present when its distributional range had some overlap with the grid cell.

#### 2.2.3 Human pressures

We obtained spatial data on the distributions of 19 human pressures which can be linked to the threats of species as listed by the IUCN red list. These included urban areas^36^, industrial areas^36^, recreational areas^36^, croplands^36^, plantation forests^37^, pastures^36^, mines^36^, wind turbines^38^, hydropower dams^39^, dams^40^, roads^41^, railways^42^, powerlines^43^, population density^44^, navigable waterways^41,42^, logging^37^, recreation potential^45^, agricultural pollutants^46^ and population pollutants^46^. All human pressure layers had information from approximately the last decade, a European or global extent, usually a spatial resolution of 1 ha or 1 km^2^, and were mostly open access (see details in Supplementary Table S1). Pressure layers with polyline or point data were turned into a raster layer with a resolution of 50 m for dams, hydro dams and wind turbines, a resolution of 100 m for navigable waterways, and a resolution of 500 m for powerlines. We turned each pressure layer into a binary raster layer (pressure = present or absent) with its original resolution. For continuous data, we used thresholds to convert each pressure layer into a binary presence-absence layer. Pressures were considered present where they have a direct footprint (e.g. presence of wind turbines, hydropower dams, dams, and powerlines), where the relevant land cover type or data class is present (e.g. urban areas, industrial areas, recreational areas, croplands, plantation forests, pastures, minefields, logging, and recreation potential), or where specific thresholds were met. These thresholds were derived from previous work and included (1) railways, population density, navigable waterways, and roads as a proxy for hunting, gathering and garbage threat^18^, (2) roads as a proxy for road threat^47,48^, and (3) environmental quality standards for population and agricultural pollution^49^. For example, we considered road pressure to be present up to a distance of 1.0 km, 1.5 km or 3.0 km on both sides of the road, depending on the taxonomic group of concern (birds and reptiles – 1.0 km threshold^47,48^, amphibians – 1.5 km threshold^48^, and mammals – 3.0 km threshold^47^). Specific details of data characteristics and thresholds of the pressure data are provided in the Supplementary Method text.

### 2.3 Linking human pressures to species threats

Each pressure layer was linked to a species-specific threat as identified and classified by the IUCN Red List^23^. Most IUCN threats (12 out of 18) could be directly represented by pressure data proxies. IUCN threats without direct pressure data (e.g. hunting, gathering, recreational activities, garbage, and water pollution threats) were indirectly represented by one or multiple pressure layers related to the IUCN threat (see Supplementary Table S1 and Supplementary Methods text). For example, Rosina *et al.* (2018)^36^ landcover data on irrigated arable land, permanently irrigated land, rice fields, vineyards, fruit trees and berry plantations, olive groves, annual crops associated with permanent crops, land principally occupied by agriculture, with significant areas of natural vegetation, and agro-forestry areas were linked to the IUCN subclass threat of annual & perennial non-timber crops (IUCN threat code 2.1). Justification for linking pressures to threats followed Allan *et al.* (2019)^18^. The pressure data allowed us to map 18 out of 45 IUCN threat subclasses that fall into 8 out of 12 IUCN threat major classes, making this the most comprehensive spatial analysis of biodiversity threats to date.

### 2.4 Spatial distribution of human pressures across European Union protected areas

We analysed the presence of each pressure in each of the EU PA grid cells, considering a pressure present when the pressure layer had some overlap with the grid cell. We calculated the area that each pressure covers within EU protected land by summing the area of all grid cells with the pressure presence. Grid cell area was calculated in ArcGIS version 10.6 using the Lambert conformal conic projection (LCC). We also calculated the co-occurrence by taking the sum of the number of pressures considered present in a grid cell.

### 2.5 Potential impacts of human pressures on threatened species

We considered a potential negative human impact to occur when three conditions were simultaneously met: 1) a pressure occurs in a grid cell, 2) a species geographic range intersects with the same grid cell, and 3) the pressure can be connected to a species-specific threat as listed by the IUCN. This assumes that human pressures are acting in a given grid cell and impacting species, and that all pressures are equally impacting a species and all species are equally impacted by a pressure. Hereafter we refer to the co-occurrence of a threat layer and the presence of a sensitive species as a ‘potential impact’^18^.

#### 2.5.1 Extent of potential human impact on sensitive threatened vertebrates in European Union protected areas

To quantify the extent of potential human impact on threatened species in EU PAs, we analysed the area of grid cells where at least one pressure is coinciding with at least one sensitive species. We also quantified the extent of potential human impact per human pressure, by analysing the total grid area where at least one sensitive species’ distribution coincides with the pressure.

Additionally, we calculated the proportion of a species’ protected range that coincides with threatening human pressures by dividing the sum of the area of grid cells where a species was considered to be potentially impacted by the sum of the area of grid cells a species was present in. We calculated this for every species across the entire EU PA network. Spatial analyses were carried out in ArcGIS version 10.6 using the Lambert conformal conic projection (LCC).

#### 2.5.2 Potential human impacts on threatened species between Natura2000 sites and non-Natura2000 sites, and between Directive and non-Directive species

We used the metric of proportion of protected species range that is potentially impacted by at least one threat to compare differences in potential impacts on species between PAs that are part of the Natura2000 and PAs outside of the Natura2000 network. We additionally classified species ranges based on whether the species is mention in Annex 1 of the EU 1979 Birds Directive^10^ or Annex 2 of the EU 1992 Habitats Directive^11^, when analysing the proportion of species’ protected ranges that are potentially impacted within and outside Natura2000 sites.

Data on the proportion of species’ protected ranges did not follow a normal distribution. Therefore, we used the Wilcoxon rank-sum test to statistically compare medians of the proportion of a potentially impacted species’ protected range in EU PAs between Natura2000 and non-Natura2000 sites, and between species mentioned in either the Birds or Habitats Directive and species not mentioned in the Directives. A p-value of < 0.05 was considered statistically significant. All statistical analyses were conducted in RStudio version 4.0.2^50^.

#### 2.5.3 Extent of potential impacts and main types of human pressures between protected area management classes

We also used the metric of proportion of species’ protected range that is potentially impacted by at least one threat to compare differences in potential impacts on species between PA management classes. We used the Kruskal–Wallis test by ranks to statistically compare medians of the potentially impacted proportion of species’ protected ranges within different PA management classes. We used the Nemenyi– Damico–Wolfe–Dunn test, from the PMCMR-package^51^, as a post hoc. A p-value of < 0.05 was considered statistically significant. All statistical analyses were conducted in RStudio version 4.0.2^50^.

## 3. Results

### 3.1 Spatial distribution of human pressures across European Union protected areas

Human pressures — i.e. activities and land use with the potential to harm threatened species — occur across 1,022,927 km^2^ (94.5%) of EU protected land (Table 1). Multiple pressures (2–16) co-occur across 978,301 km^2^ (90.3%) of EU protected land, with over half of the area (488,141 km^2^, 54.5%) being covered by > 8 co-occurring human pressures (Supplementary Figure S1). The most widespread pressures are agricultural pollution, recreation, and croplands which occur across 883,984 km^2^ (81.6%), 864,771 km^2^ (79.9%) and 797,594 km^2^ (73.7%) of EU protected land, respectively (Table 1). The spatial distribution of human pressures within EU PAs varies geographically, with hotspots of cumulative human pressure (multiple pressures co-occurring) in Luxembourg, Germany, Belgium, Poland, Czech Republic, Netherlands, Hungary, Slovenia and France (Supplementary Figure S1). Protected land with no human pressures is mostly found in Scandinavia (55,029 km^2^, 92%) (Figure 1a), and in small patches in Europe’s alpine regions (8%), including the Scottish Highlands, Alps, Pyrenees, and Carpathian Mountains (Figure 1a).

**Table 1.** Human pressures, their corresponding threat as listed by the International Union for Conservation of Nature (IUCN), number of threatened terrestrial vertebrate species sensitive to each pressure (out of 146 total), extent of pressure within European Union (EU) protected areas (PAs), and proportion of pressure extent in EU PAs where at least one species is impacted.

| Pressure | IUCN classified threat (threat code) | Number of threatened species (% total) | Area cover within EU PAs in km2 (proportion of total EU PAs) | Proportion of pressure extent impacting species |
| --- | --- | --- | --- | --- |
| Pollution (agricultural) | Agricultural & forestry effluents (9.3) | 45 (31%) | 883,984 (81.6%) | 99.99% |
| Recreation potential | Recreational activities (6.1) | 41 (28%) | 864,771 (79.9%) | 100% |
| Croplands | Annual & perennial non-timber crops (2.1) | 79 (54%) | 797,595 (73.7%) | 99.99% |
| Roads | Roads & railroads (4.1) \| Hunting & collecting terrestrial animals (5.1) \| Gathering terrestrial plants (5.2) \| Garbage & solid waste (9.4) | 85 (58%) | 794,742 (73.4%) | 100% |
| Population density | Hunting & collecting terrestrial animals (5.1) \| Gathering terrestrial plants (5.2) \| Garbage & solid waste (9.4) | 78 (53%) | 664,138 (61.3%) | 100% |
| Urban areas | Housing & urban areas (1.1) | 45 (31%) | 653,363 (60.3%) | 99.99% |
| Pastures | Livestock farming & ranching (2.3) | 47 (32%) | 524,694 (48.5%) | 99.99% |
| Pollution (urban) | Domestic & urban wastewater (9.1) | 13 (8.9%) | 470,190 (43.4%) | 89.4% |
| Logging | Logging & wood harvesting (5.3) | 21 (14%) | 412,278 (38.1%) | 95.7% |
| Railways | Roads & railroads (4.1) | 25 (17%) | 212,894 (19.7%) | 99.96% |
| Powerlines | Utility & service lines (4.2) | 11 (7.5%) | 184,751 (17.1%) | 96.3% |
| Industrial areas | Commercial & industrial areas (1.2) | 19 (13%) | 115,963 (10.7%) | 99.4% |
| Navigable waterways | Hunting & collecting terrestrial animals (5.1) \| Gathering terrestrial plants (5.2) \| Garbage & solid waste (9.4) | 78 (53%) | 105,744 (9.8%) | 100% |
| Recreational areas | Tourism & recreation areas (1.3) | 38 (26%) | 58,873 (5.4%) | 99.3% |
| Plantation forests | Wood & pulp plantations (2.2) | 23 (16%) | 27,748 (2.6%) | 76.0% |
| Mining | Mining & quarrying (3.2) | 10 (6.8%) | 18,578 (1.7%) | 17.7% |
| Dams | Dams & water management/use (7.2) | 17 (12%) | 5,808 (0.5%) | 100% |
| Windfarms | Renewable energy (3.3) | 14 (9.6%) | 5,095 (0.5%) | 100% |
| Hydro dams | Renewable energy (3.3) | 2 (1.4%) | 1,070 (0.1%) | 6.8% |
| Total |  | 139 (95%) | 1,022,927 (94.5%) | 99.3% |

**Figure 1.**
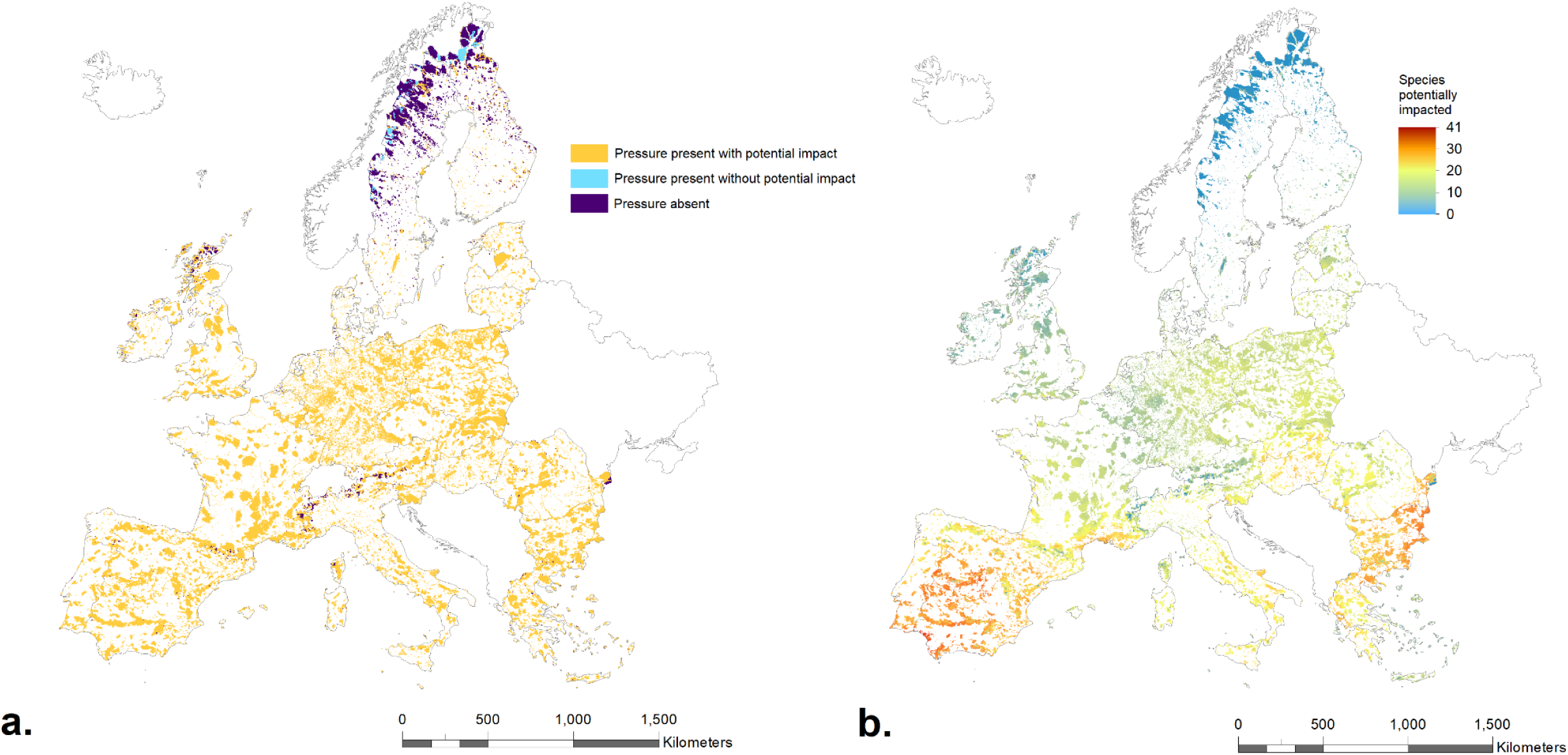
Distribution of (a) human pressures and (b) cumulative potential human impacts on threatened and near threatened vertebrates within protected areas (PAs) in the European Union. Threatened vertebrate species include amphibians, reptiles, birds and mammals (*n* = 146). PAs consists of both nationally designated PAs and PAs that are part of the Natura2000 network. Legend in (a) indicates whether a human pressure is present or absent and if it is potentially impacting at least one sensitive species within a grid cell. Legend in (b) indicates the number of sensitive species in a grid cell potentially impacted by at least one threat.

### 3.2 Extent of potential human impact on sensitive threatened vertebrates in European Union protected areas

Human pressures spatially coincided with the distribution of sensitive threatened and near threatened vertebrate species across 1,015,456 km^2^ (93.8%) of EU protected land (Figure 1a), suggesting a massive impact of human activities on species of conservation concern. Across 935,018 km^2^ (86.3%) of EU protected land, ≥ 10 species were potentially impacted (Figure 1b), and only 67,419 km^2^ (6.2%) of protected land had no species associated with threatening human pressures. Moreover, at least one species was potentially impacted across 99.3% of EU protected land where human pressure was present (Table 1). The extent of potential impacts varied considerably between different human pressure classes. We found that 15 out of the 19 pressures analysed had a potential impact on threatened species across > 95% of the EU protected land where that pressure was present, with 6 pressures (recreation potential, dams, windfarms, roads, population density and navigable waterways) potentially impacting species everywhere the pressure was present. Only mining and hydropower dam pressure were potentially impacting species across < 20% of the area where the pressure was present.

We found that 136 out of the 146 analysed species (93%) were associated with at least one threatening human pressure within their protected range (Figure 2a). Even more severely, 124 out of 146 species (85%) have > 50% of their protected range potentially impacted, and 85 out of 146 species (58%) have even > 90% of their protected range potentially impacted. Taxonomic groups differed in the proportion of a species’ protected range associated with a potential human impact, with reptiles being least affected (mean = 67.2% of protected range potentially impacted), followed by birds (mean = 79.0%), mammals (mean = 84.1%) and amphibians (mean = 85.6%) (Figure 2b–e).

**Figure 2.**
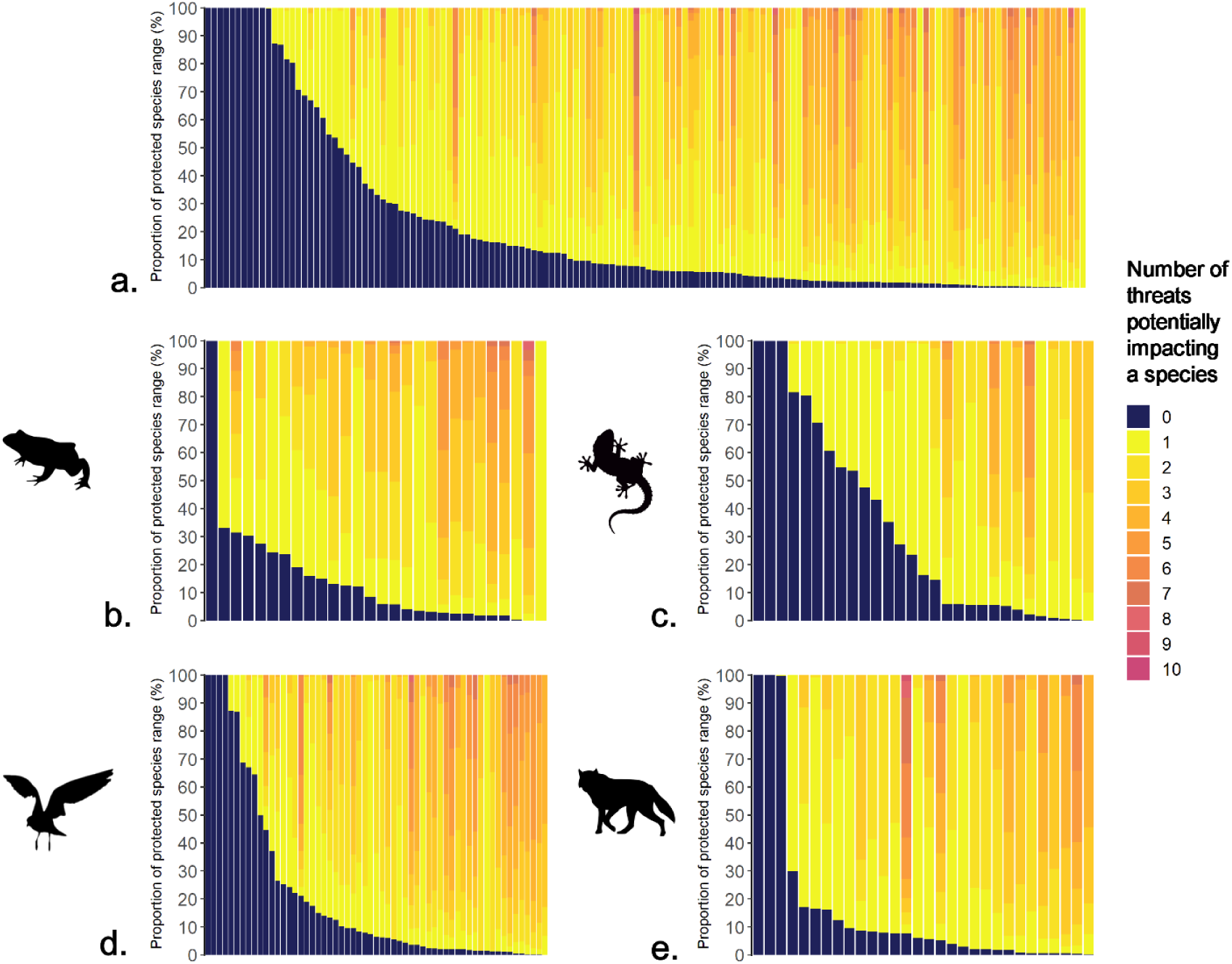
The proportion of a species range within European Union (EU) protected areas (PAs) potentially impacted by different numbers of threats. Potential impacts on species within EU PAs include distributions that overlap with regionally designated PAs and Natura2000 sites. Fractions of potential impacts within a species’ protected distribution is calculated based on a 3×3 km grid for all threatened and near threatened vertebrate species in the EU (*n* = 146; a), which includes amphibians (*n* = 28; b), reptiles (*n* = 29; c), birds (*n* = 59; d) and mammals (*n* = 30; e).

### 3.3 Potential human impacts on threatened species between Natura2000 sites and non-Natura2000 sites, and between Directive and non-Directive species

Species in Natura2000 sites were statistically less potentially impacted across their protected range than species in PAs outside of the Natura2000 network (W = 7589, *p* < 0.001) (Figure 3a). Half of all species present within Natura2000 sites were potentially impacted across ≥ 92% of their protected range, while half of all species present in PAs outside the Natura2000 network were potentially impacted across ≥ 98% of their protected range. Species mentioned in either Annex 1 of the Birds Directive or Annex 2 of the Habitats Directive were not significantly less potentially impacted than species not mentioned in either of the Directives (W = 2732, *p* = 0.76), with half of all Directive and non-Directive species being potentially impacted across ≥ 92% of their EU protected range (Figure 3b). Species mentioned in either of the Directives were also not significantly less potentially impacted than non-Directive species when focussing on ranges within Natura2000 sites or non-Natura2000 sites (W = 2713, *p* = 0.81 and W = 2585.5, *p* = 0.27 respectively) (Figure 3c, d).

**Figure 3.**
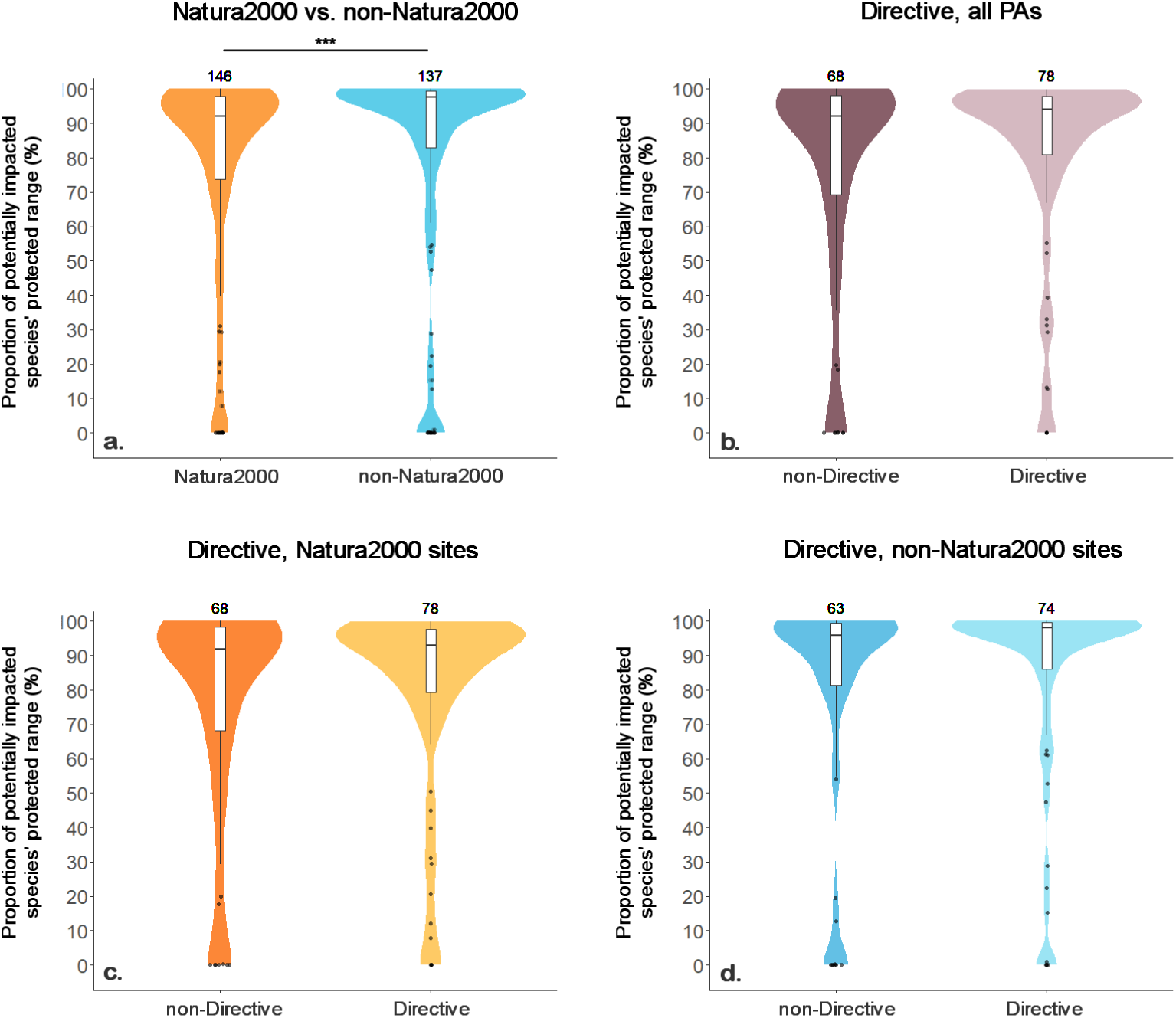
Proportion of a species European Union (EU) protected range potentially impacted by human pressures between Natura2000 and non-Natura2000 sites (a), and between species mentioned in either Annex 1 of the 1979 EU Birds Directive or Annex 2 of the 1992 EU Habitats Directive within all EA protected areas (b), within only Natura2000 sites (c) and within only sites outside the Natura2000 network (d). Species ranges of threatened and near threatened European amphibians, reptiles, birds and mammals where used. Total number of species within a category is mentioned above the corresponding bar. Proportions of a species protected range potentially impacted presented as boxplots, indicating the median, the 1^st^ and 3^rd^ quantiles, and whiskers reaching up to 1.5 times the interquartile range. Outliers are represented by black dots. The shaded areas represent violin plots of which the width correlates with the proportion of datapoints within a group. Statistical significance was calculated using the Wilcoxon rank-sum test. Statistical significance was only found between Natura2000 sites and non-Natura2000 sites, indicated by the line with asterisks (W = 7589, *p* < 0.001).

### 3.4 Extent of potential impacts and main types of human pressures between protected area management classes

The distribution of proportions of species’ protected ranges potentially impacted differed between PA management categories. Species were statistically more potentially impacted across their protected range in protected landscapes (V) than across their protected range in strict nature reserves and wilderness areas (Ia+Ib) (χ^2^= 22.9, *p* < 0.001), national parks (II) (χ^2^ = 24, *p* < 0.001), species management areas (IV) (χ^2^ = 19.73, *p* < 0.01), and protected areas with unspecified management category (N) (χ^2^ = 13.1, *p* < 0.05) (Figure 4; Supplementary Table S1). Species in protected landscapes had the highest potential impact, with half of all species present in PAs with that management category having at least 99% of their protected range potentially impacted (Figure 4). Species in strict nature reserves and wilderness areas had the lowest potential impact with half of all species present in PAs with that management category having at least 82% of their protected range potentially impacted (Figure 4).

**Figure 4.**
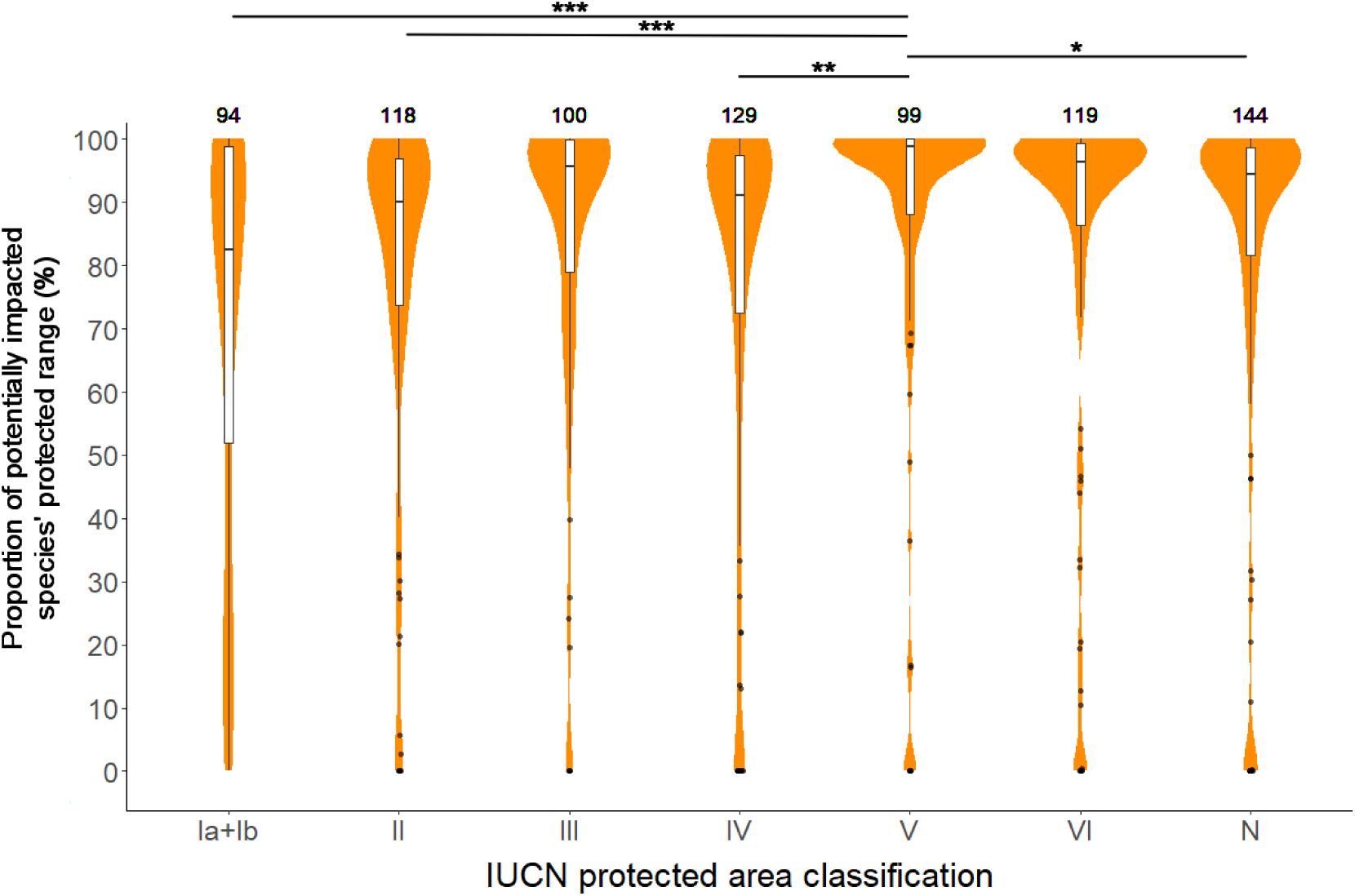
Proportion of a species European Union (EU) protected range potentially impacted by threats between protected area (PA) management classes as classified by the International Union for Conservation of Nature (IUCN) within the European Union. Species distributions of threatened and near threatened European amphibians, reptiles, birds and mammals where used. Total number of species within a class is mentioned above the corresponding bar. Proportions of a species protected range that is potentially impacted is presented as boxplots, indicating the median, the 1^st^ and 3^rd^ quantiles, and whiskers reaching up to 1.5 times the interquartile range. Outliers are represented by black dots. The shaded areas represent violin plots of which the width correlates with the proportion of datapoints within a class. Statistical significance was calculated using the Kruskal– Wallis test by ranks and the Nemenyi–Damico–Wolfe–Dunn test a post hoc test. Statistically significant differences were found between classes Ia+Ib and V, II and V, IV and V, and V and N, indicated by the lines with asterisks (χ^2^ = 22.89, *p* < 0.001; χ^2^= 23.95, *p* < 0.001; χ^2^ = 19.73, *p* < 0.01; and χ^2^ = 13.09, *p* < 0.05 respectively).

Interestingly, the type of human threats coinciding with sensitive species differed between PA management categories. Hunting & collecting of terrestrial animals potentially impacted the highest proportion of the sum of all species’ protected ranges within strictly protected areas (Ia+Ib) (15.6%), followed by recreational activities (5.9%) (Figure 5). Hunting & collecting of terrestrial animals also potentially impacted the highest proportion of the sum of all species ranges within protected area borders with less strict management (II-VI) (on average 57.1%), but was followed by non-timber crops (on average 45.9%) and agricultural effluents (on average 36.1%) (Figure 5).

**Figure 5.**
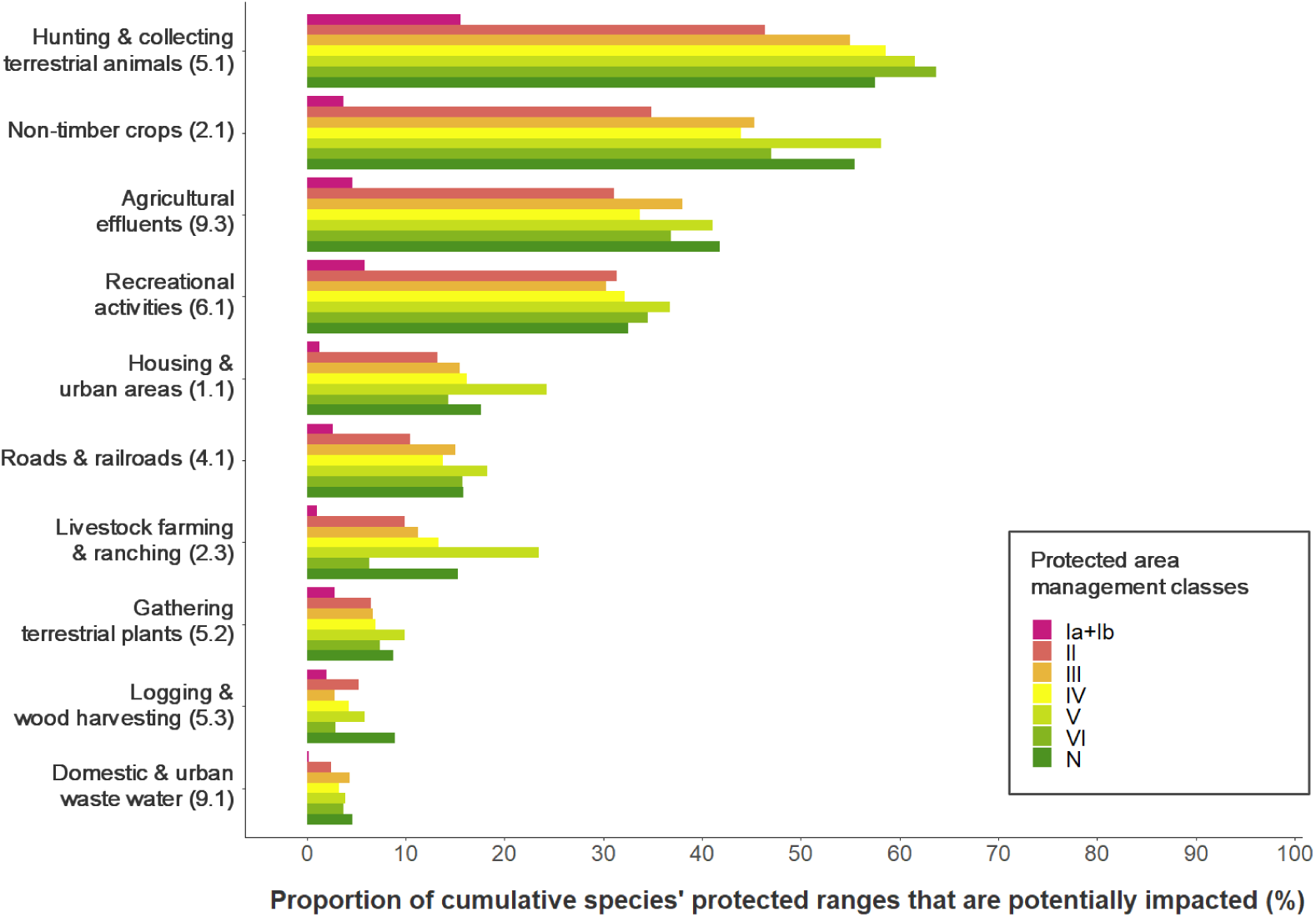
The 10 most prevalent threats as classified by the International Union for Conservation of Nature (IUCN) potentially impacting species within protected areas (PAs) of different management categories. Potential impacts on species are represented by the proportion of the sum of all species’ protected ranges within a PA management category that are coinciding with a threat. PA management classes include strict nature reserves and wilderness areas (Ia+Ib), national parks (II), species management areas (IV), natural monuments/features (V), PAs with sustainable use of natural resources (VI) and unclassified (N). Species’ protected ranges included ranges of threatened or near-threatened amphibians, reptiles, birds and mammals.

## 4. Discussion

Our analysis advances previous work that has mapped human pressures in PAs both globally and in Europe^5,7,8,52^ by accounting for the distributions of threatened vertebrates within PAs and their species-specific sensitivities to different human pressures. We therefore mapped where human pressures have a potential negative impact on species within EU PAs. Our results revealed that human activities and land uses that currently occur within EU PAs are likely harming biodiversity. We estimate that on average 84% of the protected ranges of threatened and near threatened vertebrate species in Europe are potentially impacted by one or more human pressures. This is concerning because the majority of species of European interest (listed in the EU Birds or Habitats Directive) are already in an unfavourable conservation status, and their populations are declining in PAs^53–55^. The negative impacts of human pressures on these species in places that have been set aside to protect them is particularly worrisome because it strongly increases the likelihood of their extinction^23^.

The primary aim of PAs is to effectively conserve the species living within the PA boundaries, and to ensure the long-term maintenance of healthy populations and ecosystems^56^. Human activities within protected areas should only be allowed as long as they do not negatively impact threatened species. We find that only a small part (6.2%) of the assemblages of threatened vertebrate species (i.e. all species in a grid cell) are not exposed to potential human impacts. These places are arguably the only areas of EU protected land where effective conservation is occurring. The impact of human activities on threatened species in PAs has often been overlooked in management plans, giving us a false sense of the effectiveness and sustainability of PAs in terms of biodiversity conservation^57,58^. For instance, salvage logging has been used as a management measure to reduce the spread of pest species, such as bark beetle infestations, even though it has been shown that bark beetle induced forest disturbances create valuable habitat for endangered species^59,60^.

We found that species in non-Natura2000 sites are more potentially impacted by threatening human pressures than species in Natura2000 sites. Still, 75% of all species present in Natura2000 sites are potentially impacted across 74% of their protected range. Moreover, threatened species with specific policy focus and special conservation measures (e.g. species under Annex 1 of the EU 1979 Birds Directive^10^and Annex II of the EU 1992 Habitats Directive^11^) are just as exposed to threatening human pressures as species that do not receive such attention. This result holds regardless of the PA type (Natura 2000 or not). Natura2000 sites do not necessarily focus on stricter management and less human access and use, but are rather centred around humans and nature working together^16^. The idea of such integrated management is important and in some cases species even depend on this interaction with people for survival^61^, but we show that in most cases the human activities and land uses which occur in Natura2000 sites have a potential negative impact on threatened species. A re-assessment of which activities and land uses can be allowed in particular Natura 2000 sites without negative impacts on threatened species is urgently needed to ensure that European biodiversity is effectively conserved.

We found that species in PAs with stricter IUCN management categories were less impacted than species in PAs with management more lenient towards human activities. However, even in strictly protected areas, 75% of all species present in those areas are potentially impacted across 52% of their protected range. This clearly undermines the objective of preserving species in such protected area categories^13^. This high potential impact due to human pressure within strictly protected areas could be explained by the small area (< 5 km^2^) of most strict PAs (65%) in the EU. This makes them highly accessible and precludes a well-protected core area. Most large strict EU PAs (> 350 km^2^) are only located in northern Scandinavia where species richness is low. Hence, the number of species exposed to high human pressure is mainly driven by the many small, strictly managed PAs outside northern Europe. Some of the human pressures could be mitigated through effective management efforts, such as patrolling the area for illegal hunters to mitigate hunting threat, or fencing and the building of crossing structures to reduce road threats^62,63^. However, not all pressures can be mitigated through management efforts (e.g. clear-cut logging removes all habitat of a forest species). Such activities must be prohibited from areas with sensitive species. Further work could combine our mapping of species-specific human pressures within PAs with spatial data on where mitigating management activities and conservation efforts are in place, in case such data become available. This could support policy and decision making by identifying where human pressures are currently not mitigated and thus require conservation action.

Our analysis could not include all threats for all species because spatially explicit information on many species-specific threats is missing. Pressures that were not included in this analysis were, for example, climate change^64^ and invasive species^65–67^ which both impact numerous species in European PAs. However, the impacts of these pressures will vary among species and no comprehensive spatial layers are currently available to represent species-specific threats. Some human pressure layers further underestimate the true extent of that pressure. For instance, we only included data on the impact of large dams and hydro power plants, but there are over 1 million smaller constructions that alter the flow of European rivers^68^. Our analysis also only considered the direct footprint of dams, but dams can alter the hydrology and sediment transport of river systems and therefore change entire river ecosystems^68,69^. Hence, our estimates of human impacts on threatened species in PAs are conservative and likely an underestimate of the true impact.

The accuracy of our analyses could be improved in the future if pressure datasets are developed that directly map the presence of human threats (rather than relying on proxies), and by the development of comprehensive datasets on the different mechanisms of how a pressure can impact a specific species, and their likely responses. We used the same thresholds for the presence or absence of a pressure for all species, even though each species might respond differently to the same pressure intensity. For example, different species can experience negative impacts from pollutants at different chemical concentrations. Species are also impacted by pollutants through different mechanisms, for example, they can be instantly poisoned, or chemical accumulation in body tissues can increase their vulnerability to infectious diseases or reduce their prey availability^70–72^. This will have different impacts on population dynamics and species persistence. Information could also be improved regarding species presences, since the species distribution data that we used (expert range maps) may contain commission errors (i.e. species might be falsely considered present in a given grid cell)^73^, especially when working with relatively fine grid sizes^74^. However, currently no high-resolution maps are available that represent the area of occupancy of threatened species across Europe (rather than their extent of occurrence). Moreover, the accuracy of species range maps depends on range size and expert knowledge, which varies among species, making it difficult to quantify uncertainties^73^. Once new data becomes available, our methods and analyses can be updated to assess the effect of commission errors and to improve the accuracy of the assessment for PA managers or EU member states.

## 5. Conclusion

Protected areas should be effectively managed to achieve the aim of conserving nature long-term^13^. We found that human pressures are prevalent within sensitive threatened vertebrate species ranges across 94% of EU protected land. This result is in line with other recent studies suggesting that PAs in Europe are ineffectively managed due to incomplete implementation, lack of well-informed policy decisions, and lack of resources^75–77^. A priority of the new biodiversity strategy of the European Commission therefore has to be the reduction of human pressures on habitats and species and an effective implementation of management activities to ensure the sustainable use of species and ecosystems in PAs under the new EU Nature Restoration Plan^1^. Our methodology can support the EU Member States with spatially explicit information on where human pressures affect species of policy concern across Europe, and where PA management needs to be improved to ensure the maintenance of healthy ecosystems and the services they provide. Threat free protected areas can be the cornerstone for achieving the conservation aims of the EU Biodiversity Strategy for 2030^1^.

## Supporting information

Supplementary Methods and Results

## Acknowledgements

We would like to thank Franziska Komossa, Gert-Jan Nabuurs and Alberto Pistocchi for sharing their data, making it possible to increase the detail of our spatial dataset on human pressures. W. D. Kissling acknowledges a University of Amsterdam starting grant and financial support from the Faculty Research Cluster ‘Global Ecology’.

## Literature

1. European Commission. EU Biodiversity Strategy for 2030: Bringing nature back into out lives. Communication from the Commission to the European Parliament, the Council, the European Economic and Social Committee and the Committee of the Regions. (2020); https://eur-lex.europa.eu/legal-content/EN/TXT/?qid=1590574123338&uri=CELEX:52020DC0380 (Accessed May 20, 2020)

2. CBD. Zero Draft of the Post-2020 Biodiversity Framework. (2020); https://www.cbd.int/doc/c/da8c/9e95/9e9db02aaf68c018c758ff14/wg2020-02-03-en.pdf (Accessed April, 2020)

3. Gray, C. L. et al. Local biodiversity is higher inside than outside terrestrial protected areas worldwide. Nat. Commun. 7, 12306 (2016). doi: 10.1038/ncomms12306

4. Pringle, R. M. Upgrading protected areas to conserve wild biodiversity. Nature 546, 91–99 (2017). doi: 10.1038/nature22902

5. Allan, J. R. et al. Recent increases in human pressure and forest loss threaten many Natural World Heritage Sites. Biol. Conserv. 206, 47–55 (2017). doi: 10.1016/j.biocon.2016.12.011

6. Schulze, K. et al. An assessment of threats to terrestrial protected areas. Conserv. Lett. 11, 1–10 (2018). doi: 10.1111/conl.12435

7. Geldmann, J., Manica, A., Burgess, N. D., Coad, L. & Balmford, A. A global-level assessment of the effectiveness of protected areas at resisting anthropogenic pressures. Proc. Natl. Acad. Sci. U. S. A. 116, 23209–23215 (2019). doi: 10.1073/pnas.1908221116

8. Jones, K. R. et al. One-third of global protected land is under intense human pressure. Science, 360, 788–791 (2018). doi: 10.1126/science.aap9565

9. Behre, K. E. The role of man in European vegetation history. in Vegetation History in Handbook of vegetation science, Dordrecht (eds. Huntley, B. & Webb, T.) vol. 7, 633–672 (Springer, 1988). doi: 10.1007/978-94-009-3081-0_17

10. European Economic Community. Directive 2009/147/EC of the European Parliament and the Council of 30 November 2009 on the conservation of wild birds. Off. J. L207–25 (2009).

11. European Economic Community. Council directive 92/43/EEC of 21 May 1992 on the conservation of natural habitats and of wild fauna and flora. Off. J. L2067–50 (1992).

12. EEA. Natura2000 Network Viewer. European Environment Agency (2019). Available at: https://natura2000.eea.europa.eu/#.

13. Dudley, N., Shadie, P. & Stolton, S. Guidelines for Applying Protected Area Management Categories. International Union for Conservation of Nature, Gland, Switzerland (2008).

14. Hermoso, V., Morán-Ordóñez, A., Canessa, S. & Brotons, L. Realising the potential of Natura 2000 to achieve EU conservation goals as 2020 approaches. Sci. Rep. 9, 1–10 (2019). doi: 10.1038/s41598-019-52625-4

15. Tsiafouli, M. A. et al. Human activities in Natura 2000 sites: A highly diversified conservation network. Environ. Manage. 51, 1025–1033 (2013). doi: 10.1007/s00267-013-0036-6

16. European Commission. Natura 2000. (2020). Available at: https://ec.europa.eu/environment/nature/natura2000/index_en.htm. (Accessed: June 12, 2020)

17. Martins, J. H., Camanho, A. S. & Gaspar, M. B. A review of the application of driving forces - Pressure - State - Impact - Response framework to fisheries management. Ocean Coast. Manag. 69, 273–281 (2012). doi: 10.1016/j.ocecoaman.2012.07.029

18. Allan, J. R. et al. Hotspots of human impact on threatened terrestrial vertebrates. PLoS Biol. 17, 1–18 (2019). doi: 10.1371/journal.pbio.3000158

19. Rösner, S., Mussard-Forster, E., Lorenc, T. & Müller, J. Recreation shapes a ‘landscape of fear’ for a threatened forest bird species in Central Europe. Landsc. Ecol. 29, 55–66 (2014). doi: 10.1007/s10980-013-9964-z

20. Garrido, M. & Pérez-Mellado, V. Human pressure, parasitism and body condition in an insular population of a Mediterranean lizard. Eur. J. Wildl. Res. 61, 617–621 (2015). doi: 10.1007/s10344-015-0915-7

21. Suraci, J. P., Clinchy, M., Zanette, L. Y. & Wilmers, C. C. Fear of humans as apex predators has landscape-scale impacts from mountain lions to mice. Ecol. Lett. 22, 1578–1586 (2019). doi: 10.1111/ele.13344

22. Berger, J. Fear, human shields and the redistribution of prey and predators in protected areas. Biol. Lett. 3, 620–623 (2007). doi: 10.1098/rsbl.2007.0415

23. IUCN. The IUCN Red List of Threatened Species. Version 2020-1 (2020). Available at: https://www.iucnredlist.org. (Accessed: June 12, 2020)

24. Jung, K. & Threlfall, C. G. Trait-dependent tolerance of bats to urbanization: A global meta-analysis. Proc. R. Soc. B., 285, 20181222 (2018). doi: 10.1098/rspb.2018.1222

25. Venter, O. et al. Sixteen years of change in the global terrestrial human footprint and implications for biodiversity conservation. Nat. Commun. 7, 1–11 (2016). doi: 10.1038/ncomms12558

26. O’Bryan, C. J. et al. Intense human pressure is widespread across terrestrial vertebrate ranges. Glob. Ecol. Conserv. 21, e00882 (2020). doi: 10.1016/j.gecco.2019.e00882

27. Loos, J. et al. Low-intensity agricultural landscapes in Transylvania support high butterfly diversity: Implications for conservation. PLoS One 9, e103256 (2014). doi: 10.1371/journal.pone.0103256

28. Doxa, A. et al. Low-intensity agriculture increases farmland bird abundances in France. J. Appl. Ecol. 47, 1348–1356 (2010). doi: 10.1111/j.1365-2664.2010.01869.x

29. Oldekop, J. A., Holmes, G., Harris, W. E., Evans, K. L. & Evaluaci, U. A global assessment of the social and conservation outcomes of protected areas. Conservation Biology, 30, 133–141 (2015). doi: 10.1111/cobi.12568

30. Thorn, S. et al. Impacts of salvage logging on biodiversity: a meta-analysis. J. Appl. Ecol., 55, 279–289 (2018). doi: 10.1111/1365-2664.12945

31. UN Environment World Conservation Monitoring Centre. World Database on Protected Areas. International Union for Conservation of Nature (2020). Available at: https://www.protectedplanet.net/ (Accessed January 27, 2020).

32. European Environment Agency. Natura 2000 data - the European network of protected sites. (2018). Available at: https://www.eea.europa.eu/data-and-maps/data/natura-11#tab-gis-data. (Accessed January 27, 2020)

33. Leroux, S. J. et al. Global protected areas and IUCN designations: Do the categories match the conditions? Biol. Conserv. 143, 609–616 (2010). doi: 10.1016/j.biocon.2009.11.018

34. BirdLife. Bird species distribution maps of the world. BirdLife International and Handbook of the Birds of the World (2019). Available at: http://datazone.birdlife.org/species/requestdis. (Accessed February 5, 2020)

35. Roll, U. et al. The global distribution of tetrapods reveals a need for targeted reptile conservation. Nat. Ecol. Evol. 1, 1677–1682 (2017). doi: 10.1038/s41559-017-0332-2

36. Rosina, K. et al. Increasing the detail of European land use/cover data by combining heterogeneous data sets. Int. J. Digit. Earth 13, 602–626 (2018). doi: 10.1080/17538947.2018.1550119

37. Nabuurs, G. et al. Next-generation information to support a sustainable course for European forests. Nat. Sustain. 2, 815–818 (2019). doi: 10.1038/s41893-019-0374-3

38. Rehbein, J. A. et al. Renewable energy development threatens many globally important biodiversity areas. Glob. Chang. Biol. 26, 3040–3051 (2020). doi: 10.1111/gcb.15067

39. Lehner, B. et al. High-resolution mapping of the world’s reservoirs and dams for sustainable river-flow management. Front. Ecol. Environ. 9, 494–502 (2011). doi: 10.1890/100125

40. Mulligan, M., van Soesbergen, A. & Sáenz, L. GOODD, a global dataset of more than 38,000 georeferenced dams. Sci. Data 7, 31 (2020). doi: 10.1038/s41597-020-0362-5

41. Eurogeographics. EuroGlobalMap 2019 (2019). Available at: https://eurogeographics.org/maps-for-europe/open-data/topographic-data/ (Accessed March 1, 2020)

42. Venter, O. et al. Global terrestrial Human Footprint maps for 1993 and 2009. Sci. Data 3, 1–10 (2016). doi: 10.1038/sdata.2016.67

43. Gotovtsev, M. OpenStreetMap data in shapefile format. (2016). Available at: osm2shp.ru (Accessed February 25, 2020)

44. Schiavina, M., Freire, S. & MacManus, K. GHS-POP R2019A - GHS population grid multitemporal (1975-1990-2000-2015). European Commission, Joint Research Centre (JRC) (2019). [Dataset] doi: 10.2905/0C6B9751-A71F-4062-830B-43C9F432370F

45. Komossa, F., van der Zanden, E. H., Schulp, C. J. E. & Verburg, P. H. Mapping landscape potential for outdoor recreation using different archetypical recreation user groups in the European Union. Ecol. Indic. 85, 105–116 (2018). doi: 10.1016/j.ecolind.2017.10.015

46. Pistocchi, A., Dorati, C., Aloe, A., Ginebreda, A. & Marcé’, R. River pollution by priority chemical substances under the Water Framework Directive: A provisional pan-European assessment. Sci. Total Environ. 662, 434–445 (2019). doi: 10.1016/j.scitotenv.2018.12.354

47. Torres, A., Jaeger, J. A. G. & Alonso, J. C. Assessing large-scale wildlife responses to human infrastructure development. PNAS 113, 8472–8477 (2016). doi: 10.1073/pnas.1522488113

48. Brehme, C. S., Hathaway, S. A. & Fisher, R. N. An objective road risk assessment method for multiple species: ranking 166 reptiles and amphibians in California. Landsc. Ecol. 33, 911–935 (2018). doi: 10.1007/s10980-018-0640-1

49. European Economic Commission. Directive 2008/105/EC of the European Parliament and of the Council of 16 December 2008 on environmental quality standards in the field of water policy, amending and subsequently repealing Council Directives 82/176/EEC, 83/513/EEC, 84/156/EEC, 84/491/EEC, 86/280/EEC and amending Directive 2000/60/EC of the European Parliament and of the Council. Official Journal of the European Union, OJ L 348, p. 84–97 (2008).

50. RStudio Team. RStudio: Integrated Development for R. RStudio, Inc., Boston, MA. (2016).

51. Pohlert, T. The Pairwise Multiple Comparison of Mean Ranks Package (PMCMR). R Package version 4.3. (2014). Available at: https://CRAN.R-project.org/package=PMCMR

52. Geldmann, J., Joppa, L. N. & Burgess, N. D. Mapping Change in Human Pressure Globally on Land and within Protected Areas. Conserv. Biol. 28, 1604–1616 (2014). doi: 10.1111/cobi.12332

53. European Environment Agency. Biodiversity briefing. (2015). Available at: https://www.eea.europa.eu/soer/2015/europe/biodiversity. (Accessed: 16th June 2020)

54. Palacín, C. & Alonso, J. C. Failure of EU Biodiversity Strategy in Mediterranean farmland protected areas. J. Nat. Conserv. 42, 62–66 (2018). doi: 10.1016/j.jnc.2018.02.008

55. Sergio, F. et al. Protected areas under pressure: Decline, redistribution, local eradication and projected extinction of a threatened predator, the red kite, in Doñana National Park, Spain. Endanger. Species Res. 38, 189–204 (2019). doi: 10.3354/ESR00946

56. Oliver, T. H. et al. Biodiversity and Resilience of Ecosystem Functions. Trends Ecol. Evol. 30, 673–684 (2015). doi: 10.1016/j.tree.2015.08.009

57. Meinig, H. U. & Boye, P. A review of negative impact factors threatening mammal populations in Germany. Folia Zool. 58, 279–290 (2009).

58. European Commission. Commission notice: Managing Natura 2000 Sites. The provisions of Article 6 of the ‘Habitats’ Directive 92/43/EEC. Office for Official Publications of the European Communities, Luxembourg. (2018).

59. Müller, J. et al. Increasing disturbance demands new policies to conserve intact forest. Conserv. Lett. 12, 1–7 (2019). doi: 10.1111/conl.12449

60. Beudert, B. et al. Bark beetles increase biodiversity while maintaining drinking water quality. Conserv. Lett. 8, 272–281 (2015). doi: 10.1111/conl.12153

61. Moreira, F., Pinto, M. J., Henriques, I. & Marques, T. The importance of low-intensive farming systems for fauna, flora and habitats protected under the european ‘Birds’ and ‘Habitats’ Directives: is agriculture essential for preserving biodiversity in the mediterranean region? in Trends in Biodiversity Research in Nova Science Publishers, New York (eds. Burks, A. R.) 117–145 (2005).

62. Van der Ree, R. et al. Effects of roads and traffic on wildlife populations and landscape function: road ecology is moving towards larger scales. Ecol. Soc. 16, 48 (2011).

63. Moore, J. F. et al. Are ranger patrols effective in reducing poaching-related threats within protected areas? J. Appl. Ecol. 55, 99–107 (2018). doi: 10.1111/1365-2664.12965

64. Araújo, M. B., Alagador, D., Cabeza, M., Nogués-Bravo, D. & Thuiller, W. Climate change threatens European conservation areas. Ecol. Lett. 14, 484–492 (2011). doi: 10.1111/j.1461-0248.2011.01610.x

65. European Environment Agency. The impacts of invasive alien species in Europe. EEA Technical report, No 16/2012. (2012).

66. Gallardo, B. et al. Protected areas offer refuge from invasive species spreading under climate change. Glob. Chang. Biol. 23, 5331–5343 (2017). doi: 10.1111/gcb.13798

67. Guerra, C., Baquero, R. A., Gutiérrez-Arellano, D. & Nicola, G. G. Is the Natura 2000 network effective to prevent the biological invasions? Glob. Ecol. Conserv. 16, (2018). doi: 10.1016/j.gecco.2018.e00497

68. Gough, P., Fernández Garrido, P. & Van Herk, J. Dam Removal. A viable solution for the future of our European rivers. Dam Removal Europe (2018).

69. Lange, K. et al. Small hydropower goes unchecked. Front. Ecol. Environ. 17, 256–258 (2019). doi: 10.1002/fee.2049

70. BirdLife International. Species factsheet: Tetrax tetrax. (2020). Available at: http://datazone.birdlife.org/species/factsheet/little-bustard-tetrax-tetrax/text. (Accessed: 16th April 2020)

71. McCoy, K. A. & Peralta, A. L. Pesticides could alter amphibian skin microbiomes and the effects of Batrachochytrium dendrobatidis. Front. Microbiol. 9, 1–5 (2018). doi: 10.3389/fmicb.2018.00748

72. Ali, F. et al. A review of the effects of some selected pyrethroids and related agrochemicals on aquatic vertebrate biodiversity. Canadian Journal of Pure and Applied Sciences 5, 1455–1464 (2011).

73. Rondinini, C., Wilson, K. A., Boitani, L., Grantham, H. & Possingham, H. P. Tradeoffs of different types of species occurrence data for use in systematic conservation planning. Ecol. Lett. 9, 1136–1145 (2006). doi: 10.1111/j.1461-0248.2006.00970.x

74. Di Marco, M., Watson, J. E. M., Possingham, H. P. & Venter, O. Limitations and trade-offs in the use of species distribution maps for protected area planning. J. Appl. Ecol. 54, 402–411 (2017). doi: 10.1111/1365-2664.12771

75. Getzner, M., Jungmeirer, M. & Pfleger, B. Evaluating management effectiveness of national parks as a contribution to good governance and social learning. in Protected area management, IntechOpen (eds. Sladonja, B.) (2012). doi: 10.5772/50092. Available from: https://www.intechopen.com/books/protected-area-management/evaluating-management-effectiveness-of-national-parks-as-a-contribution-to-good-governance-and-socia

76. Kati, V. et al. The challenge of implementing the European network of protected areas Natura 2000. Conserv. Biol. 29, 260–270 (2015). doi: 10.1111/cobi.12366

77. Hermoso, V., Morán-Ordóñez, A., Canessa, S. & Brotons, L. Four ideas to boost EU conservation policy as 2020 nears. Environ. Res. Lett. 14, 101001 (2019). doi: 10.1088/1748-9326/ab48cc

