## Supplementary Methods and Results for "Threatened terrestrial vertebrates are exposed to human pressures across 94% of Europe’s protected land"

*Data availability of IUCN classified species threats within the European Union*

**Table S1. Information on use and availability of datasets that can represent IUCN classified major classes and sub-classes of threats to species. Information about the datasets includes whether the pressure data directly (D) or indirectly (I) represent the threat class, year of information content, spatial resolution and extent of the dataset as well as availability of the dataset.**

| | IUCN threat code | IUCN threat class | Indirect (I) or direct (D) pressure proxy (type of data when indirect proxy) | Year | Resolution | Extent | Open Access | Reference |
|---|---|---|---|---|---|---|---|---|
| 1. Residential & commercial development | 1.1 | Housing & urban areas | D | 2012 | 1ha | Europe | Yes | Rosina *et al.* (2018)[1] |
| | 1.2 | Commercial & industrial areas | D | 2012 | 1ha | Europe | Yes | Rosina *et al.* (2018)[1] |
| | 1.3 | Tourism & recreation areas | D | 2012 | 1ha | Europe | Yes | Rosina *et al.* (2018)[1] |
| 2. Agriculture & aquaculture | 2.1 | Annual & perennial non-timber crops* | D | 2012 | 1ha | Europe | Yes | Rosina et al. (2018)[1] |
| | 2.2 | Wood & pulp plantations* | D | 2000 - 2012 | 1km2 | Europe | Upon request | Nabuurs et al. (2019)[2] |
| | 2.3 | Livestock farming & ranching* | D | 2012 | 1ha | Europe | Yes | Rosina et al. (2018)[1] |
| | 2.4 | Marine & freshwater aquaculture* | - | - | - | - | - | - |

| | | | | | | | | |
|---|---|---|---|---|---|---|---|---|
| 3. Energy production & mining | 3.1 | Oil & gas drilling | - | - | - | - | - | - |
| | 3.2 | Mining & quarrying | D | 2012 | 1ha | Europe | Yes | Rosina et al. (2018)[1] |
| | 3.3 | Renewable energy | D | 2018 | Point data | Global | Yes | Lehner *et al.* (2011)[3] |
| | | | D | 2017 | Point data | Global | Upon request | Rehbein *et al.* (2020)[4] |
| 4. Transportation & service corridors | 4.1 | Roads & railroads | D | 2019 | Polyline data | Europe | Yes | Eurogeographics (2019)[5] |
| | | | D | 2009 | 1 km2 | Global | Yes | Venter et al. (2016)[6] |
| | 4.2 | Utility & service lines | D | 2016 | Polyline data | Europe | Yes | Gotovtsev (2018)[7] |
| | 4.3 | Shipping lanes | - | - | - | - | - | - |
| | 4.4 | Flight paths | - | - | - | - | - | - |
| 5. Biological resource use | 5.1 | Hunting & collecting terrestrial animals* | I (Population density) | 2015 | 250m | Global | Yes | Schiavina *et al.* (2019)[8] |
| | | | I (Paved roads + navigable waterways) | 2019 | Polyline data | Europe | Yes | Eurogeographics (2019)[5] |
| | | | I (Navigable waterways) | 2009 | 1 km2 | Global | Yes | Venter et al. (2016)[6] |
| | 5.2 | Gathering terrestrial plants* | I (Population density) | 2015 | 250m | Global | Yes | Schiavina *et al.* (2019)[8] |
| | | | I (Paved roads + navigable waterways) | 2019 | Polyline data | Europe | Yes | Eurogeographics (2019)[5] |
| | | | I (Navigable waterways) | 2009 | 1 km2 | Global | Yes | Venter et al. (2016)[6] |
| | 5.3 | Logging & wood harvesting* | D | 2000 - 2012 | 1km2 | Europe | Upon request | Nabuurs *et al.* (2019)[2] |
| | 5.4 | Fishing & harvesting aquatic resources* | - | - | - | - | - | - |

| | | | | | | | | |
|---|---|---|---|---|---|---|---|---|
| 6. Human intrusion & disturbance | 6.1 | Recreational activities | I (Outdoor recreation potential) | 2012-2018 | 1km2 | EU | Upon request | Komossa *et al.* (2018)[9] |
| | 6.2 | War, civil unrest & military exercises | - | - | - | - | - | - |
| | 6.3 | Work & other activities | - | - | - | - | - | - |
| 7. Natural system modifications | 7.1 | Fire & fire suppression* | - | - | - | - | - | - |
| | 7.2 | Dams & water management/use* | D | 2020 | Point data | Global | Yes | Mulligan *et al.* (2020)[10] |
| | 7.3 | Other ecosystem modifications | - | - | - | - | - | - |
| 8. Invasive & other problematic species, genes & diseases* | n/a | n/a | - | - | - | - | - | - |
| 9. Pollution | 9.1 | Domestic & urban waste water* | I | 2000-2012 | Based on river catchments[11] | EU | Upon request | Pistocchi *et al.* (2019)[12] |
| | 9.2 | Industrial & military effluents* | - | - | - | - | - | - |
| | 9.3 | Agricultural & forestry effluents* | I | 2000-2012 | Based on river catchments[11] | EU | Upon request | Pistocchi et al. (2019)[12] |
| | 9.4 | Garbage & solid waste | I (Population density) | 2015 | 250m | Global | Yes | Schiavina *et al.* (2019)[8] |
| | | | I (Paved roads + navigable waterways) | 2019 | Polyline data | Europe | Yes | Eurogeographics (2019)[5] |
| | | | I (Navigable waterways) | 2009 | 1 km2 | Global | Yes | Venter et al. (2016)[6] |
| | 9.5 | Air-borne pollutants* | - | - | - | - | - | - |
| | 9.6 | Excess energy* | - | - | - | - | - | - |

| | | | | | | | | |
|---|---|---|---|---|---|---|---|---|
| 10. Geological events | n/a | n/a | - | - | - | - | - | - |
| 11. Climate change & severe weather | n/a | n/a | - | - | - | - | - | - |
| 12. Other options | n/a | n/a | - | - | - | - | - | - |

* Contains sub-classes

*Linking species threats as classified in the IUCN Red List with human pressures (IUCN threat code)*

*Housing & urban areas (1.1)*

Under this category IUCN gives the following examples: 'urban areas, suburbs, villages, ranchettes, vacation homes, shopping areas, offices, schools, hospitals, birds flying into windows, land reclamation or expanding human habitation that causes habitat degradation in riverine, estuary and coastal areas, etc'. These examples are directly captured by six landcover classes within the Rosina *et al.* (2018)[1] dataset. These landcover classes include four intensities of urban fabric ranging from >50% to <10% built-up, public facilities and construction sites. These classes were all converted into a binary pressure layer (pressure present or absent). Pressure was considered present when the landcover contained one of the previously described classes.

*Commercial & industrial areas (1.2)*

Under this category IUCN gives the following examples: 'military bases, factories, stand-alone shopping centres, office parks, power plants, train yards, ship yards, airports, landfills, etc.'. These examples are directly captured by seven landcover classes within the Rosina *et al.* (2018)[1] dataset. These landcover classes include production facilities, commercial/service facilities, major railway stations, port areas, airport areas, airport terminals and dump sites. These classes were all converted into a binary pressure layer (pressure present or absent). Pressure was considered present when the landcover contained one of the previously described classes.

*Tourism & recreation areas (1.3)*

Under this category IUCN gives the following examples: 'ski areas, golf courses, resorts, cricket fields, county parks, afghan goat polo fields, campgrounds, coastal and estuarine tourist resorts, etc'. These examples are directly captured by two landcover classes within the Rosina *et al.* (2018)[1] dataset. These landcover classes include sport and leisure facilities and sport, leisure and touristic built-up. These classes were all converted into a binary pressure layer (pressure present or absent). Pressure was considered present when the landcover contained one of the previously described classes.

*Annual & perennial non-timber crops (2.1)*

IUCN considers 'crops planted for food, fodder, fibre, fuel, or other uses' to fall under this category. Scale of the farming activity can range from shifting agriculture, small-holder farming and agro-industry farming. This is directly captured by nine landcover classes within the Rosina *et al.* (2018)[1] dataset. These landcover classes irrigated arable land, permanently irrigated land, rice fields, vineyards, fruit trees and berry plantations, olive groves, annual crops associated with permanent crops, land principally occupied by agriculture, with significant areas of natural vegetation and agro-forestry areas. These classes were all converted into a binary pressure layer (pressure present or absent). Pressure was considered present when the landcover contained one of the previously described classes.

*Wood & pulp plantations (2.2)*

IUCN considers 'stands of trees planted for timber or fibre outside of natural forests, often with non-native species' to fall under this category. Scale of the plantations can range from small-holder plantations to agro-industry plantations. The probable presence of these forest plantations is captured under the very intensive forest management category from Nabuurs *et al.* (2019)[2] as this category represents forest patches under a short rotation cycle which are in most cases fast growing tree species, for example eucalypt farms in Portugal. These classes were all converted into a binary pressure layer (pressure present or absent). Pressure was considered present when the management type contained one of the previously described classes.

*Livestock farming & ranching (2.3)*

IUCN considers 'domestic terrestrial animals raised in one location on farmed or nonlocal resources (farming); also domestic or semi-domesticated animals allowed to roam in the wild and supported by natural habitats (ranching)' to fall under this category. The scale of this threat varies from nomadic grazing and small-holder or agro-industry grazing, ranching or farming. This is directly captured by the pasture landcover class within the Rosina *et al.* (2018)[1] dataset. This class was converted into a binary pressure layer (pressure present or absent). Pressure was considered present when the landcover contained the previously described classes.

*Mining & quarrying (3.2)*

Under this category IUCN gives the following examples: 'coal strip mines, alluvial gold panning, gold mines, rock quarries, sand/salt mines, coral mining, deep sea nodules, guano harvesting, dredging outside of shipping lanes, etc.'. This is partially captured by the mineral extraction site landcover class within the Rosina *et al.* (2018)[1] dataset. This class was converted into a binary pressure layer (pressure present or absent). Pressure was considered present when the landcover contained the previously described classes.

*Renewable energy (3.3)*

Under this category IUCN gives the following examples: 'geothermal power production, solar farms, wind farms (including birds flying into windmills), tidal farms, etc.'. Some of these examples are directly captured, which include hydropower dams and wind turbines. These are both captured by the data on hydropower dams locations from Lehner *et al.* (2011)[3] and by the data on wind turbine locations form Rehbein *et al.* (2020)[4]. Most species used in this analysis are threatened by wind turbines, followed by a small amount of species being threatened by hydropower dams. Because all species used in this study are threatened by only one type of renewable energy form this threat is divided into a hydropower dam and wind turbine subclass. Only threats regarding hydropower dams and wind turbines were further used in the analysis with a separate pressure layer for each of the two classes. One renewable energy threat was excluded from a species, because it was threatened by tidal farms and could not be represented by the pressure data. Both hydropower dam and wind turbine locations were converted into a binary pressure layer (pressure present or absent). Pressure was considered present within a 50×50 m square raster around the hydropower dam or wind turbine locations. A 50×50 m square was chosen to limit the chance of the pressure being present within two neighbouring planning units.

*Roads & railroads (4.1)*

Under this category IUCN gives the following examples: 'highways, secondary roads, primitive roads, logging roads, bridges & causeways, road kill, fencing associated with roads, freight/passenger/mining railroads, etc.'. This is directly captured by the paved road data from Eurogeographics (2019)[5] and by the data on railroads from Venter *et al.* (2016)[6]. Species are affected by this threat through road mortality, ecosystem degradations or disturbances. For this reason we chose to exclusively use paved roads, because cars can drive on higher speeds and surrounding habitats experience more change by the paving of the road than dirt roads[13]. Pressure was considered present up to 1 km either side of the road for birds and amphibians, 1.5 km either side of the road for reptiles, and 3 km either side of the road for mammals or up to 500m either side of the railway for every animal type. These railroad threshold choices were made by following Allan *et al.* (2019)[14]. Road threshold choices for birds and mammals were made based on changes in mean species abundances studied by Torres *et al.* (2016)[13]. Road threshold choices for amphibians and reptiles were made by taking the mid value of population movement distances of species of which this distance had a medium to high confidence in its estimation in the study of Brehme *et al.* (2018)[15].

*Utility & service lines (4.2)*

Under this category IUCN gives the following examples: ‘electrical & phone wires, aqueducts, oil & gas pipelines, electrocution of wildlife, etc.’. This is directly captured by the powerline data from Gotoytsev (2018)[7] based on the OpenStreetMap data. Categories that were included in this analysis were 'line', 'line;minor_line', 'low_line', 'minor', 'minor_line', 'minor_line; line', 'minor_line;line' or 'wire'. Pressure was considered present up to 500m of the powerline location. The strongest impact of electrical wires on species is through electrocution and direct contact (death). However, we chose a 500m threshold due to computing power when converting the polyline data to a raster file, and to include more diffuse impacts (e.g. avoidance of areas with powerlines).

*Hunting & collecting terrestrial animals (5.1)*

Under this category IUCN gives the following examples: ‘bushmeat hunting, trophy hunting, beaver trapping, butterfly collecting, honey or bird nest hunting, pest control often impacts non-targeted species, hunter's dogs may chase after and kill other non-target species during a hunt, loss of a species' prey base due to over-harvesting by humans of their prey, wolf control, pest control, persecution of snakes because of superstition, etc.’. As there is no data covering the direct presence of hunting, we used indirect proxies to represent this layer. Following Allan *et al.* (2019)[14] we used roads and navigable waterways together with human population density to represent the hunting threat. We used paved road data from Eurogeographics (2019)[5], navigable waterway data from Venter *et al.* (2016)[6] and Eurogeographics (2019)[5], and data on population density from Schiavina *et al.* (2019)[8]. Pressure was considered present up to 3 km either side of the road, 1.5 km either side of the navigable waterway or with a population density of >1 person/km$^2$.

*Gathering terrestrial plants (5.2)*

Under this category IUCN gives the following examples: 'wild mushroom collection, forage for stall fed animals, orchid collection, rattan harvesting, other plants accidentally removed/killed as a result of methods/approach used to harvest a target species, etc.'. As there is no data covering the direct presence of gathering, we used proxies to represent this layer. Following the methods from Allan *et al.* (2019)[14] we used roads and navigable waterways together with human population density to represent the gathering threat. We used paved road data from Eurogeographics (2019)[5], navigable waterway data from Venter *et al.* (2016)[6] and Eurogeographics (2019)[5], and data on population density from Schiavina *et al.* (2019)[8]. Pressure was considered present up to 3 km either side of the road, 1.5 km either side of the navigable waterway or with a population density of >1 person/km$^2$.

*Logging & wood harvesting (5.3)*

IUCN considers 'harvesting trees and other woody vegetation for timber, fibre, or fuel' to fall under this category. Both intentional and unintentional use and both small and large scale logging is considered within this threat category. The probable presence of logging is captured under the very intensive, intensive and multifunctional forest management category from Nabuurs *et al.* (2019)[2]. These classes were all converted into a binary pressure layer (pressure present or absent). Pressure was considered present when the management type contained one of the previously described classes.

*Recreational activities (6.1)*

Under this category IUCN gives the following examples: 'off-road vehicles, motorboats, motorcycles, jet-skis, snowmobiles, ultralight planes, dive boats, whale watching, mountain bikes, hikers, cross-country skiers, hang gliders, birdwatchers, scuba divers, pets brought into recreation areas, temporary campsites, caving, rock-climbing, etc.'. There is no direct data on recreation activities in Europe, however Komossa *et al.*

(2018)[9] did calculate the outdoor recreation potential of the EU area based on landscape preferences of different recreation groups. We used the recreation potentials of all recreation groups as an indicator of recreational activity threat on species. Pressure was considered present in highly accessible areas with medium to high recreation potential, as recreational activities mostly cause species stresses and changes in behaviour that can have negative effect on species when disturbed frequently[16].

*Dams & water management/use (7.2) – subcategories Large Dams (7.2.10) and Dams (size unknown) (7.2.11)*

IUCN considers ‘changing water flow patterns from their natural range of variation either deliberately or as a result of other activities’ to fall under this category. There is no data available for water management but there is data available to represent the subcategories focussed on dams (7.2.10 and 7.2.11) This is captured by the dam location data from Mulligan *et al.* (2020)[10]. Pressure was considered present within a 50×50 m square raster around the dam locations. A 50×50 m square was chosen to limit the chance of the pressure being present within two neighbouring planning units.

*Domestic & urban waste water (9.1)*

Under this category IUCN gives the following examples: ‘discharge from municipal waste treatment plants, leaking septic systems, untreated sewage, outhouses, oil or sediment from roads, fertilizers and pesticides from lawns and golf-courses, road salt, etc.’. This is partially captured by chemical pollution data from Pistocchi *et al.* (2019)[12] which included chemical concentrations of population pollutants per river catchment[11]. As impacts caused by pollution differ per species and per chemical we decided to use the maximum allowed concentration of priority pollutants under the Water Framework Directive[17]. Pressure was considered present when a domestic/urban related chemical would succeed the maximum allowed concentration. The following chemicals were included in creating this pressure layer: tributyltin, lead, nonylphenol, nickel, mercury, hexachlorobutadiene, fluoranthene, chloroalkanes (C10-C13), and anthracene.

*Agricultural & forestry effluents (9.1)*

Under this category IUCN gives the following examples: ‘nutrient loading from fertilizer run-off, manure from feedlots, nutrients from aquaculture, herbicide run-off from orchards, etc.'. This is partially captured by chemical pollution data from Pistocchi *et al.* (2019)[12] which included chemical concentrations of agricultural pollutants per river catchment[11]. As impacts caused by pollution differ per species and per chemical we decided to use the maximum allowed concentration of priority pollutants under the Water Framework Directive[17]. Pressure was considered present when a domestic/urban related chemical would succeed the maximum allowed concentration. The following chemicals were included in creating this pressure layer: chlorpyrifos, cypermethrin, chlorfenvinphos, cadmium, bifenox, alachlor, terbutryn, pentacholophenol, isoproturon, hexachloro-cyclohexane, hexachlorobenzene, heptachlor, endosulfan, diuron, and dichlorvos.

*Garbage & solid waste (9.4)*

Under this category IUCN gives the following examples: 'municipal waste, litter from cars, flotsam & jetsam from recreational boats, waste that entangles wildlife, construction debris, etc.'. This is indirectly captured by road data from Eurogeographics (2019)[5], navigable waterway data from Venter *et al.* (2016)[6] and Eurogeographics (2019)[5], and data on population density from Schiavina *et al.* (2019)[8]. Gathering terrestrial plant pressure was  considered present up to 3 km either side of the road, 1.5 km either side of the navigable waterway or with a population density of >1 person/km$^2$.

### Supplementary Results

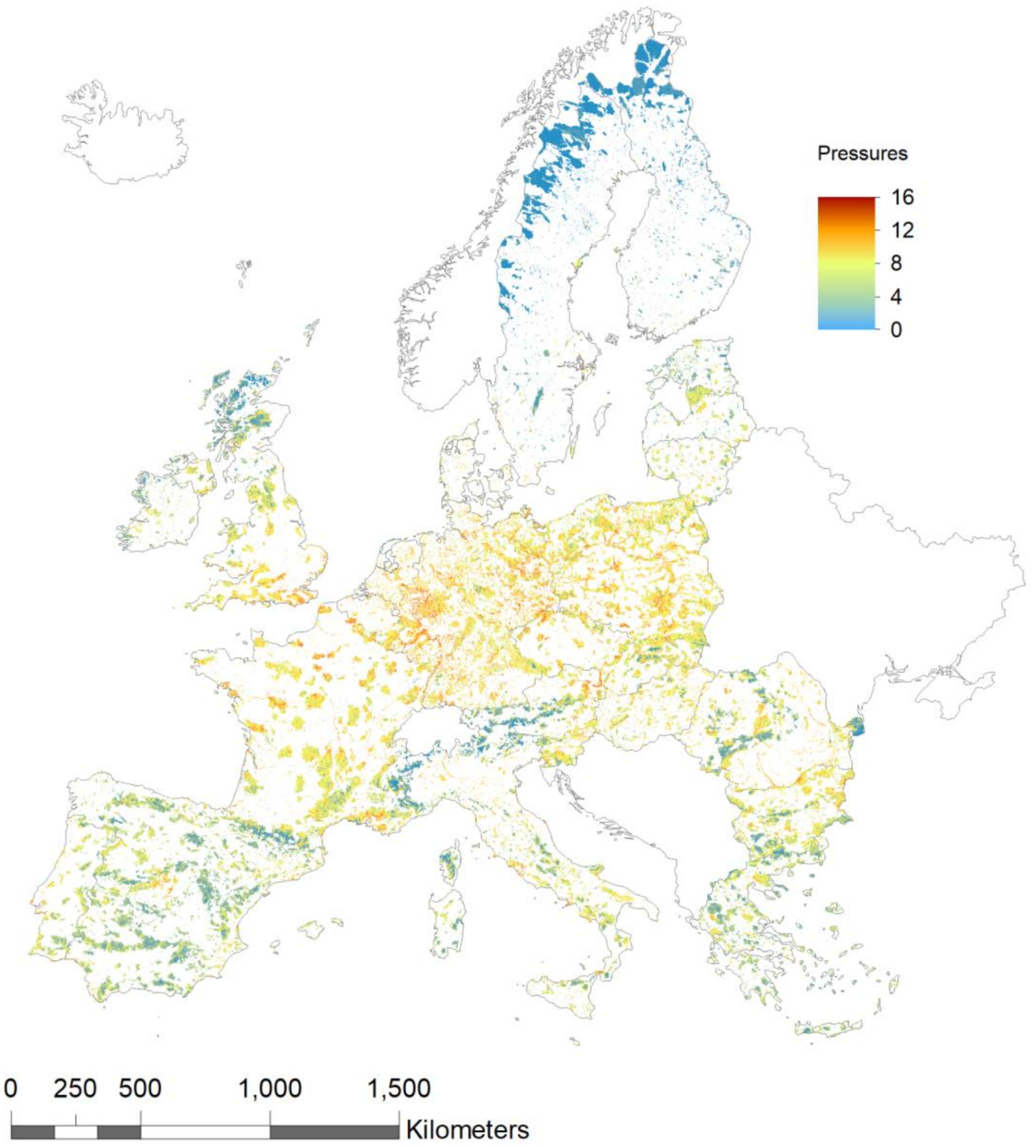


**Figure S1. Number of human pressures present within protected areas (PAs) in the European Union.** PAs consists of both nationally designated PAs and PAs that are part of Natura2000.

**Table S1. Pairwise comparisons of proportion of impacted species range between protected area management classes using Nemenyi–Damico–Wolfe–Dunn test with Chi-squared ($\chi^2$) approximation.** Per International Union for Conservation of Nature (IUCN) classified management class the median proportion of impacted protected species range is given as well as the sample size. A p-value < 0.05 was considered statistically significant and is shown in bold red.

| | | | | Management class | | | | | |
|---|---|---|---|---|---|---|---|---|---|
| **Management class** | **Median** | **n** | | **II** | **III** | **IV** | **V** | **VI** | **N** |
| **Ia+Ib** | 82.5 | 94 | Test statistic ($\chi^2$) | 0.0254 | 8.35 | 0.497 | **22.9** | 7.79 | 2.67 |
| | | | p-value | 1.0 | 0.21 | 1.0 | **0.00083** | 0.25 | 0.85 |
| **II** | 90.1 | 118 | Test statistic ($\chi^2$) | - | 8.37 | 0.334 | **24** | 7.82 | 2.46 |
| | | | p-value | | 0.21 | 1.0 | **0.00053** | 0.25 | 0.87 |
| **III** | 95.8 | 100 | Test statistic ($\chi^2$) | | - | 5.75 | 3.73 | 0.0488 | 2.33 |
| | | | p-value | | | 0.45 | 0.71 | 1.0 | 0.89 |
| **IV** | 91 | 129 | Test statistic ($\chi^2$) | | | - | **19.73** | 5.19 | 0.998 |
| | | | p-value | | | | **0.0031** | 0.52 | 0.98 |
| **V** | 94.4 | 144 | Test statistic ($\chi^2$) | | | | - | 1.85 | **13.1** |
| | | | p-value | | | | | 0.93 | **0.042** |
| **VI** | 98.9 | 99 | Test statistic ($\chi^2$) | | | | | - | 4.99 |
| | | | p-value | | | | | | 0.55 |
| **N** | 86.4 | 119 | Test statistic ($\chi^2$) | | | | | | - |
| | | | p-value | | | | | | |

**Literature - Supplementary**


1. Rosina, K. *et al.* Increasing the detail of European land use/cover data by combining heterogeneous data sets. *Int. J. Digit. Earth* **13**, 602–626 (2018). doi:10.1080/17538947.2018.1550119
2. Nabuurs, G. *et al.* Next-generation information to support a sustainable course for European forests. *Nat. Sustain.* **2**, 815-818 (2019). doi:10.1038/s41893-019-0374-3
3. Lehner, B. *et al.* High-resolution mapping of the world's reservoirs and dams for sustainable river-flow management. *Front. Ecol. Environ.* **9**, 494–502 (2011). doi:10.1890/100125
4. Rehbein, J. A. *et al.* Renewable energy development threatens many globally important biodiversity areas. *Glob. Chang. Biol.* **26**, 3040–3051 (2020). doi:10.1111/gcb.15067
5. Eurogeographics. EuroGlobalMap 2019 (2019). Available at: https://eurogeographics.org/maps-for-europe/open-data/topographic-data/ (Accessed March 1, 2020)
6. Venter, O. *et al.* Global terrestrial Human Footprint maps for 1993 and 2009. *Sci. Data* **3**, 1–10 (2016). doi:10.1038/sdata.2016.67
7. Gotovtsev, M. *OpenStreetMap data in shapefile format*. (2016). Available at: osm2shp.ru (Accessed February 25, 2020)
8. Schiavina, M., Freire, S. & MacManus, K. GHS-POP R2019A - GHS population grid multitemporal (1975-1990-2000-2015). *European Commission, Joint Research Centre (JRC)* (2019). [Dataset] doi:10.2905/0C6B9751-A71F-4062-830B-43C9F432370F
9. Komossa, F., van der Zanden, E. H., Schulp, C. J. E. & Verburg, P. H. Mapping landscape potential for outdoor recreation using different archetypical recreation user groups in the European Union. *Ecol. Indic.* **85**, 105–116 (2018). doi:10.1016/j.ecolind.2017.10.015
10. Mulligan, M., van Soesbergen, A. & Sáenz, L. GOODD, a global dataset of more than 38,000 georeferenced dams. *Sci. Data* **7**, 1–8 (2020). doi:10.1038/s41597-020-0362-5
11. Vogt, J. *et al.* A pan-European river and catchment database. *EC-JRC report EUR 22920 EN*, *Luxembourg* (2007). doi:10.2788/35907
12. Pistocchi, A., Dorati, C., Aloe, A., Ginebreda, A. & Marcé', R. River pollution by priority chemical substances under the Water Framework Directive: A provisional pan-European assessment. *Sci. Total Environ.* **662**, 434–445 (2019). doi:10.1016/j.scitotenv.2018.12.354
13. Torres, A., Jaeger, J. A. G. & Alonso, J. C. Assessing large-scale wildlife responses to human infrastructure development. *PNAS* **113**, 8472–8477 (2016). doi:10.1073/pnas.1522488113
14. Allan, J. R. *et al.* Hotspots of human impact on threatened terrestrial vertebrates. *PLoS Biol.* **17**, 1–18 (2019). doi:10.1371/journal.pbio.3000158

15. Brehme, C. S., Hathaway, S. A. & Fisher, R. N. An objective road risk assessment method for multiple species: ranking 166 reptiles and amphibians in California. *Landsc. Ecol.* **33**, 911–935 (2018). doi:10.1007/s10980-018-0640-1

16. Larson, C. L., Reed, S. E., Merenlender, A. M. & Crooks, K. R. A meta-analysis of recreation effects on vertebrate species richness and abundance. *Conservation Science and Practice* **1**, 1–9 (2019). doi:10.1111/csp2.93

17. European Economic Commission. Directive 2008/105/EC of the European Parliament and of the Council of 16 December 2008 on environmental quality standards in the field of water policy, amending and subsequently repealing Council Directives 82/176/EEC, 83/513/EEC, 84/156/EEC, 84/491/EEC, 86/280/EEC and amending Directive 2000/60/EC of the European Parliament and of the Council. *Official Journal of the European Union,* OJ L 348, p. 84–97 (2008).
